# Effect of Neck Muscle Strength and Passive Cervical Viscoelasticity on Regional Brain and Axonal Tissue Response

**DOI:** 10.64898/2026.10.27.762371

**Authors:** Hossein Bahreinizad, Gustavo M. Paulon, Michael S. Jaffee, Lakiesha N. Williams, Suman K. Chowdhury

**Affiliations:** Department of Industrial and Systems Engineering, Herbert Wertheim College of Engineering, University of Florida, Gainesville, FL, USA; Department of Neurology, College of Medicine, University of Florida, Gainesville, FL, USA; J. Crayton Pruitt Family Department of Biomedical Engineering, Herbert Wertheim College of Engineering, University of Florida, Gainesville, FL, USA

**Keywords:** traumatic brain injury, finite element modeling, neck muscle strength, cervical viscoelasticity, axonal strain, intracranial pressure, head-neck biomechanics, diffusion tensor imaging

## Abstract

This study evaluated how neck muscle strength and passive cervical viscoelasticity influence global brain motion, intracranial pressure, and regional predicted axonal strain during rapid head loading. An MRI-derived human head-neck finite element model with 5000 diffusion-derived axonal tracts represented by 90,692 embedded beam elements was evaluated using a 4 × 4 factorial design, with four levels each of muscle strength and passive ligament/disc viscoelasticity. The 16 cervical configurations were simulated under a 15 g acceleration-driven inertial load and a 7000 N frontal impact, resulting in 32 simulations. Increasing muscle strength from S25 to S95 at the reference passive-viscoelasticity level reduced mean regional 95th-percentile predicted axonal strain by 22% under inertial loading and 18% under frontal loading, with regional reductions ranging from 10% to 58% and 7% to 31%, respectively. Increasing passive cervical viscoelasticity from D25 to D95 at the reference strength level produced larger reductions in mean regional axonal strain of 35% and 33%, with regional reductions ranging from 14% to 70% and 16% to 56%. Passive viscoelasticity also reduced brain center-of-mass angular acceleration by 16% and 13%, compared with reductions of only 1% and 2% with increased muscle strength. Regional intracranial pressure decreased by 9%–21% under inertial loading and 5%–12% under frontal loading with increased passive viscoelasticity, whereas greater muscle strength reduced axonal strain without a corresponding reduction in pressure. The effects of the two cervical factors also depended on one another: under inertial loading, the mean axonal-strain reduction from increasing strength increased from 21% at D25 to 28% at D95, while the corresponding passive-viscoelasticity effect increased from 32% at S25 to 38% at S95. These findings show that cervical mechanical properties can substantially alter regional brain and axonal response, with passive cervical viscoelasticity producing larger effects than muscle strength across the ranges evaluated.

## 1. Introduction

Traumatic brain injury (TBI) remains a major global health problem, with approximately 69 million cases estimated worldwide each year (Dewan et al., 2019). Mechanical loading of the head produces translational and rotational motion, intracranial pressure gradients, brain motion relative to the skull, and deformation of brain tissue. These internal responses are difficult to measure directly during impact in living humans. Finite element (FE) models therefore provide an important tool for relating external loading to the mechanical response of the brain. Experimental and computational studies have been used to identify the biomechanical factors that determine how external head loading produces motion and deformation within the brain (Kleiven, 2007; Alshareef et al., 2018; Zhou et al., 2018). While much of this work has focused on head kinematics, brain-skull motion, intracranial pressure, and brain deformation (Kleiven, 2007; Giordano & Kleiven, 2014; Alshareef et al., 2018), more recent studies have increasingly considered the neck as part of the coupled mechanical system that influences head and brain response (Jin et al., 2017; Kuo et al., 2019; Mortensen et al., 2020; Cournoyer et al., 2021).

One component of this cervical mechanical state is neck muscle strength, which represents the available force-generating capacity of the cervical musculature. Neck strength varies substantially among individuals. Catenaccio et al. (2017) measured 157 healthy young adults and reported strength ranges of 38–383 N in men and 15–223 N in women, while Vasavada et al. (2008) also reported substantial interindividual differences. Strength can also be increased through training (Geary et al., 2014). However, evidence that greater strength consistently reduces head loading or concussion risk remains mixed. Eckner et al. (2014) found inverse relationships between isometric neck strength and changes in head linear and angular velocity during controlled impulsive loading, with correlation coefficients ranging from 0.417 to 0.657. In contrast, Mihalik et al. (2011) found that greater cervical strength did not consistently reduce head-impact severity in youth ice hockey players and reported higher head-impact severity among athletes with greater upper-trapezius strength. Schmidt et al. (2014) similarly found no protective effect of greater cervical muscle strength or size on football head-impact severity, while Garrett et al. (2023) reported only small, nonsignificant pooled relationships between greater strength and lower concussion risk. Computational studies also show that cervical muscle effects depend on more than force capacity alone. Jin et al. (2017) found that both the level and strategy of neck muscle activation affected predicted mild-TBI response, indicating that how the muscles are recruited can be as important as their available force capacity. Mortensen et al. (2020) found that doubling isometric strength reduced HIC by approximately 6–10%, BrIC by 0.5–6%, and HIP by 4–9%, while posture and impact conditions produced larger changes. Together, these mixed findings suggest that muscle force-generating capacity alone may not fully describe the influence of the neck on head and brain mechanics. Passive cervical mechanics represent another component of the head-neck system that may influence the response to rapid loading.

Passive cervical tissues also influence head-neck mechanics. Cervical ligaments exhibit nonlinear viscoelastic behavior that depends on strain magnitude and loading rate (Lucas et al., 2008; Troyer & Puttlitz, 2011, 2012). Intervertebral discs also exhibit frequency-dependent stiffness and damping, although much of the available dynamic human disc data comes from lumbar specimens (Izambert et al., 2003). At the whole-neck level, Kuo et al. (2019) showed that the relative contributions of active muscle and passive ligaments changed with impact severity; during simulated football impacts, the active-muscle contribution during acceleration fell below 20%, while passive ligaments became increasingly important. Dirisala et al. (2012) likewise found that introducing damping into a simplified FE neck reduced later peaks in intracranial pressure, shear stress, and shear strain. These findings indicate that passive cervical viscoelasticity should be considered separately from muscle force-generating capacity.

Importantly, the influence of cervical mechanics is not limited to gross head motion. Alvarez et al. (2014) varied neck muscle tonus during simulated pedestrian impacts and reported changes in brain strain of approximately 1–14% across most conditions. Cournoyer et al. (2021) similarly found that changes in neck stiffness altered both head kinematics and maximum principal strain, with the magnitude and direction of the effect depending on impact velocity, location, and striking mass. Global motion also does not necessarily reflect local brain deformation. Zhou et al. (2018) found no significant relationship between peak brain-skull relative motion and peak brain strain and no significant relationship between FE validation scores based on these two response types. Cervical effects on TBI mechanics should therefore be evaluated using intracranial measures in addition to global head or brain motion.

Among these intracranial responses, axonal deformation is particularly relevant because white matter is organized into directionally oriented pathways. Bain and Meaney (2000) experimentally linked dynamic white-matter stretch with morphological and electrophysiological axonal injury. FE studies subsequently showed that deformation resolved along fiber directions can differ substantially from conventional maximum principal strain (Giordano & Kleiven, 2014; Ji et al., 2015; Zhao et al., 2016). Ji et al. (2015), for example, reported that at a strain threshold of 0.18, fiber-oriented strain identified 3.2% of white matter above the threshold compared with 29.8% using maximum principal strain. More recent approaches have represented reconstructed white-matter pathways explicitly as embedded one-dimensional elements, allowing deformation to be evaluated directly along the modeled tracts (Wu et al., 2019). Hajiaghamemar et al. (2020) further showed in an animal model that embedded-tract metrics predicted histologically identified traumatic axonal injury with accuracies of 73–90% and generally outperformed corresponding tissue-level measures. These findings support evaluating deformation along reconstructed axonal pathways rather than relying only on continuum brain strain.

Despite these advances, the combined effects of muscle force-generating capacity and passive cervical viscoelasticity on regional brain and axonal mechanics remain poorly characterized. Experimental studies have shown that cervical strength and neuromuscular state can influence head kinematics, although their relationship with concussion risk remains inconsistent (Eckner et al., 2014; Schmidt et al., 2014; Garrett et al., 2023). Computational studies have shown that muscle properties and passive cervical mechanics can alter head-neck response and continuum brain deformation (Mortensen et al., 2020; Kuo et al., 2019; Dirisala et al., 2012; Alvarez et al., 2014; Cournoyer et al., 2021). In parallel, brain FE models have progressed from fiber-oriented strain measures to direct representation of reconstructed axonal pathways (Giordano & Kleiven, 2014; Zhao et al., 2016; Wu et al., 2019). However, muscle strength, passive cervical mechanics, and axonal deformation have largely been studied separately. The combined effects of graded muscle strength and passive ligament-and-disc viscoelasticity on regional intracranial pressure and deformation along explicitly modeled axonal pathways have not been systematically evaluated in a coupled head-neck model. It therefore remains unclear whether the effect of muscle strength depends on the passive mechanical state of the neck, whether the effect of passive cervical viscoelasticity depends on muscle strength, and whether these relationships change between acceleration-driven inertial loading and contact-driven frontal impact.

The objective of this study was therefore to quantify the effects of neck muscle strength and passive cervical viscoelasticity on brain center-of-mass kinematics, brain-skull relative motion, regional intracranial pressure, and regional predicted axonal strain using an MRI-derived head-neck FE model with embedded diffusion-derived axonal tracts. Muscle strength and passive cervical viscoelasticity were each evaluated at four literature-informed levels in a complete 4 × 4 parameter grid, producing 16 cervical configurations under each of two mechanically distinct loading conditions: a 15 g acceleration-driven inertial condition and a contact-driven frontal impact with a peak force of 7000 N. We hypothesized that both cervical factors would alter brain response, that passive cervical viscoelasticity would produce the larger effect during rapid loading, and that the magnitude of each factor effect would depend on the level of the other factor and on the loading condition.

## 2. Materials and Methods

### 2.1 MRI-derived head-neck finite element model

A previously developed and validated MRI-derived human head-neck FE model was used as the baseline anatomical framework (Bahreinizad et al., 2025). The model was developed from MRI data from a healthy 42-year-old male participant who was 176 cm tall and weighed 106 kg. It includes the scalp, skull, dura mater, pia mater, cerebrospinal fluid, gray matter, white matter, C1-C7 cervical vertebrae, intervertebral discs, 42 cervical muscles, and 14 cervical ligament structures. The complete model contains more than 1.46 million elements. Gray matter, white matter, and cerebrospinal fluid are represented with hexahedral elements, while the cervical vertebrae, discs, and other complex structures are represented with solid elements appropriate to their geometry. Cervical muscles and ligaments are represented using one-dimensional elements.

The baseline head-neck model was previously evaluated against whole head-neck kinematics from the Naval Biodynamics Laboratory data, cervical intervertebral-disc deformation from Ito et al. (2005), and brain displacement from the sonomicrometry experiments of Alshareef et al. (2018) (Bahreinizad et al., 2025).

### 2.2 DTI-derived axonal tract model

Diffusion tensor imaging (DTI) data were acquired using the updated UF diffusion MRI protocol with 128 diffusion sampling directions, b-values up to 2000 s/mm^2^, and a spatial resolution of 2 × 2 × 2 mm^3^. The diffusion data were processed in DSI Studio. Motion and eddy-current corrections were applied before deterministic whole-brain tractography. Tractography was performed using an FA threshold of 0.20, an angular threshold of 60°, a 2 mm step size, and a stopping criterion of 5000 streamlines.

The reconstructed streamlines were registered to the FE brain geometry before conversion to beam elements. Streamline coordinates were exported from DSI Studio and aligned with the FE brain geometry in MATLAB using an iterative closest-point (ICP) procedure. The ICP algorithm iteratively calculated the rigid transformation that minimized the distance between the tractography data and the corresponding FE brain geometry, and the resulting transformation was applied to all streamline coordinates. The registered streamlines were then imported into ANSA and visually inspected relative to the FE brain mesh. Residual local misalignments were manually corrected in ANSA to ensure that the tract trajectories were properly positioned within the brain domain. After registration and manual correction, the tract curves were discretized into 2 mm segments and converted into 90,692 one-dimensional tension-only beam elements representing 5000 reconstructed tracts.

**Figure 1.**
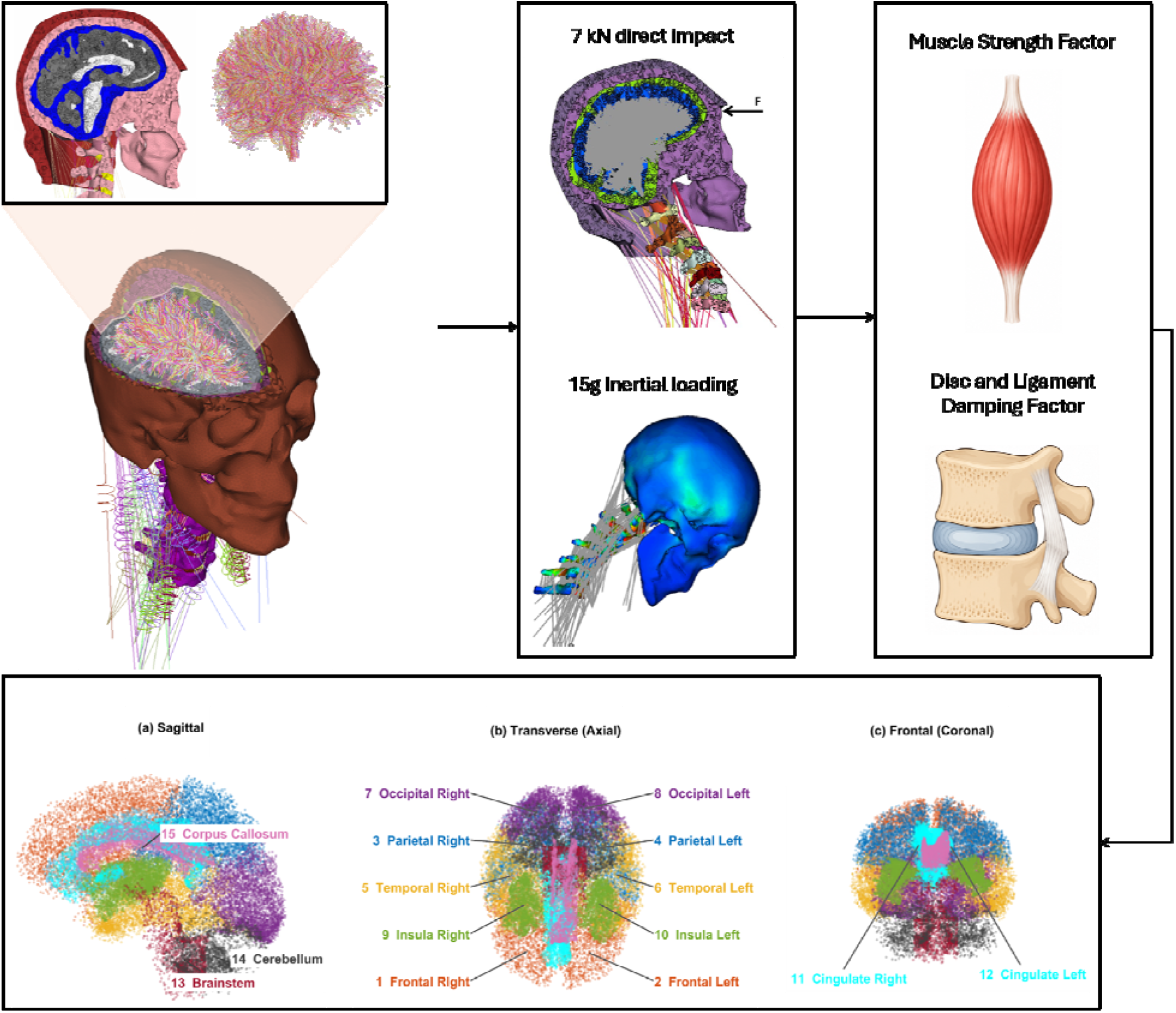
Overview of the modeling framework used in this study. MRI-derived head-neck anatomy and diffusion-derived white-matter tracts were integrated to construct the head-neck finite element model with embedded axonal pathways. Cervical muscle strength and passive ligament/disc viscoelasticity were varied as the two neck-mechanical factors, and the model was subjected to the prescribed loading conditions. Regional brain responses were evaluated using 15 atlas-based regions, shown in sagittal, transverse (axial), and frontal (coronal) views.

FA values sampled along each streamline were averaged to obtain a tract-specific FA value, and this value was assigned to the beam elements belonging to that tract. The axonal beam network was embedded within the brain solid domain using the LS-DYNA constrained beam-in-solid formulation. Beam density was assigned a negligible value so that the embedded network contributed negligible additional mass to the brain model.

### 2.3 Brain ground-substance and axonal material models

The brain was represented using the ground-substance and axonal material framework of Wu et al. (2019). Brain solid elements represented the ground substance, while the embedded tension-only beam elements represented the axonal contribution. The same ground-substance formulation was used for gray matter and for the solid component of white matter. White matter additionally contained the embedded DTI-derived axonal beam elements.

The strain-energy density of the brain ground substance was defined as

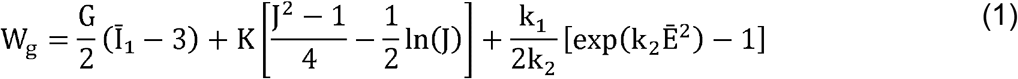

where the isotropic ground-substance strain-like term was

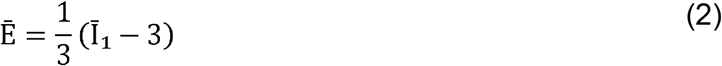

The strain-energy density of the embedded axonal component was

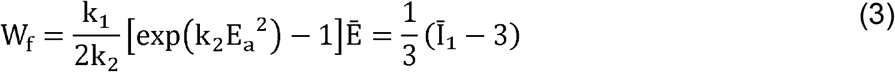

with the axonal strain-like term

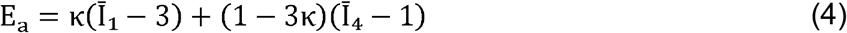

and the fiber pseudo-invariant

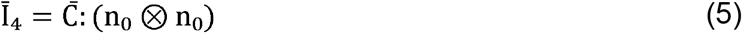

The fiber dispersion parameter was calculated from the tract-specific FA value as

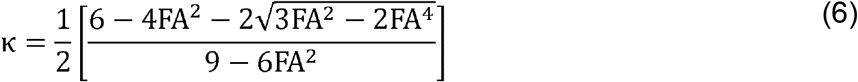

The time-dependent deviatoric response was represented using a reduced relaxation function

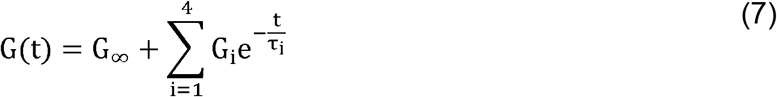

The ground-substance parameters were G = 27.5 kPa, K = 2.19 GPa, k_1_ = 203 kPa, and k_2_ → 0. The axonal parameters were k_1_ = 900 kPa and k_2_ → 0, with κ determined separately for each tract from FA. The reduced relaxation coefficients were G_1_ = 0.8087, G_2_ = 0.1005, G_3_ = 0.0470, G_4_ = 0.0133, and G∞ = 0.0305. The corresponding relaxation times were τ_1_ =0.01 ms, τ_2_ = 0.23 ms, τ_3_ = 5 ms, and τ_4_ = 200 ms (Wu et al., 2019). For k_2_ → 0, the axonal exponential energy term reduces to

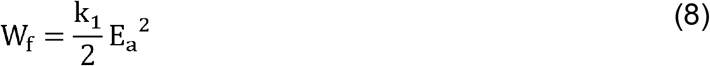

### 2.4 Brain-response validation

The updated brain model was evaluated using the same Alshareef et al. (2018) sonomicrometry case used in the previous head-neck model study. Alshareef et al. applied dynamic pure rotational pulses and measured three-dimensional brain displacement with implanted sonomicrometry crystals. Following Bahreinizad et al. (2025), the present validation reproduces the 40 rad/s coronal pulse with a 60 ms duration in the lateral direction. Cervical muscle activation is set to 0% to represent the postmortem condition, and the head-neck model is scaled by 0.92 to match the reported 1.265 kg brain mass. Brain displacement is extracted at the same three reported receiver locations used in the previous validation and compared with the experimental X-, Y-, and Z-direction displacement histories.

### 2.5 Cervical muscle activation and muscle-strength variation

The model contains 21 bilateral cervical muscle groups, resulting in 42 modeled muscles. Muscle force was represented using a Hill-type formulation in which total muscle force was the sum of active and passive components:

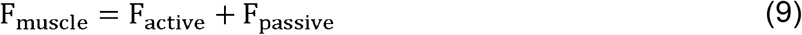

The active contractile force was

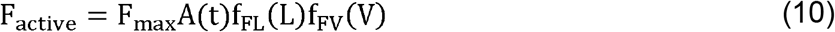

where Fmax is maximum isometric force, *A*(*t*)is muscle activation, *f*_*FL*_ is the normalized force-length relation, and *f*_*Fv*_ is the normalized force-velocity relation. The passive force was defined as

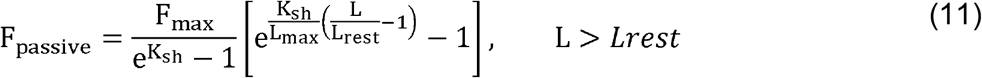

and F_passive_ = 0 for L ≤ L_rest_. This formulation follows the Hill-type cervical muscle implementation described by Panzer et al. (2011) and used in the previously published head-neck model (Bahreinizad et al., 2025).

Muscle activation patterns were estimated using a modified HYOID musculoskeletal model in OpenSim. The HYOID muscle set was modified to correspond directly to the 21 bilateral muscle groups represented in the FE model. Following the approach of Mortensen et al. (2018), the head-neck model was maintained in a neutral posture and a resistance load was applied to generate the required cervical muscle effort. OpenSim static optimization was used to determine the muscle activation pattern by minimizing the sum of squared muscle activations while satisfying mechanical equilibrium. Low-capacity reserve coordinate actuators were included to provide any residual generalized forces or moments required to obtain an equilibrium solution. The reserve actuators were assigned low capacities so that the applied load was carried primarily by the modeled muscles rather than by the reserve actuators. Reserve-actuator forces and moments were used only in the OpenSim optimization and were not transferred to the FE model. The resulting muscle activation values were mapped to the corresponding 42 Hill-type muscles in the FE model and were held fixed across all muscle-strength conditions. Therefore, the strength analysis changed maximum force-generating capacity while preserving the same muscle recruitment pattern.

Reference maximum isometric force values followed the cervical muscle model of Panzer et al. (2011), which used a maximum isometric muscle stress of 0.5 MPa. Interindividual strength variability was based on the male flexion and extension measurements reported by Catenaccio et al. (2017), with Vasavada et al. (2008) providing independent support for substantial human neck-strength variability. A sagittal coefficient of variation of 0.283 was calculated from the reported strength data and combined with standard-normal quantiles to define nominal S25, S50, S75, and S95 whole-neck strength levels. The resulting multiplicative factors were 0.809, 1.000, 1.191, and 1.466. The same factor was applied to the maximum isometric force of every modeled cervical muscle:

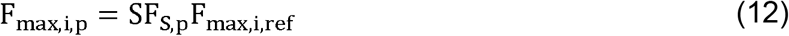

Muscle activation, force-length behavior, force-velocity behavior, resting length, and passive muscle properties were unchanged across the strength conditions.

**Table 1.**
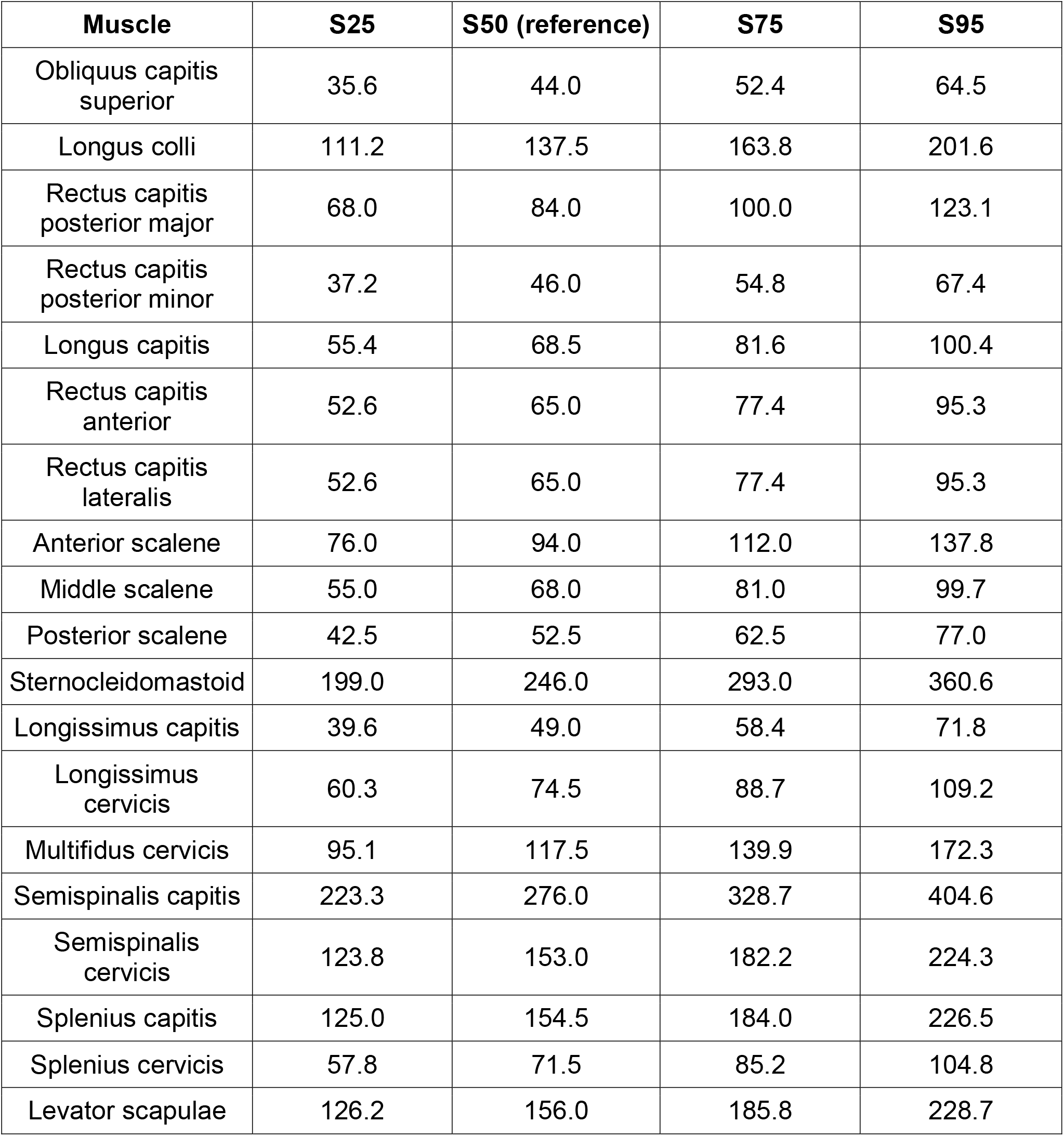
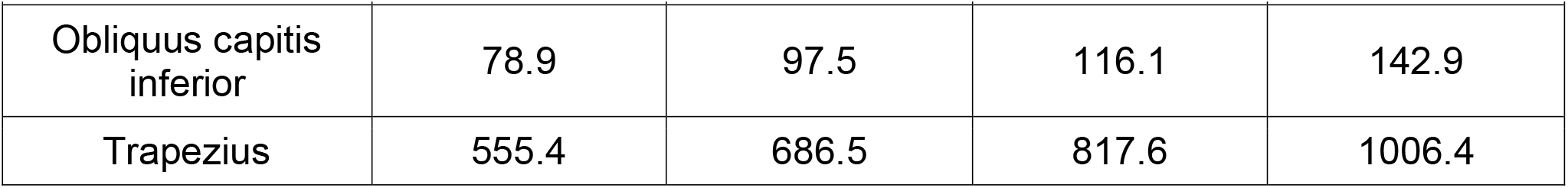
Maximum isometric force (N) values used for the muscle-strength conditions.

### 2.6 Cervical ligament viscoelasticity

The cervical ligament sensitivity model used the nonlinear viscoelastic formulation reported by Troyer and Puttlitz (2012). The time-dependent strain-dependent modulus was

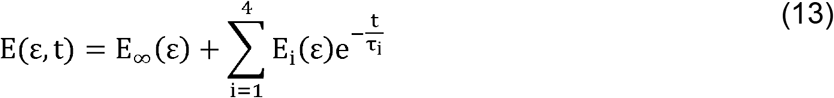

where *E*_∞_ (ε) is the equilibrium strain-dependent modulus and *E*_1_ (ε) through *E*_4_ (ε) are the transient strain-dependent components. The relaxation times were *τ*_1_ = 0.1*s, τ*_2_ = 1*s, τ*_3_ = 10*s*, and *τ*_4_ = 100*s* and were held fixed across all ligament sensitivity conditions (Troyer & Puttlitz, 2012).

Variability in the transient ligament response was estimated from the fast-rate human cervical ligament data of Lucas et al. (2008). A transient coefficient of variation of approximately 0.20 was calculated from the reported relaxation-response data across the anterior longitudinal ligament, posterior longitudinal ligament, and ligamentum flavum. This value was used as the sensitivity magnitude. The resulting ligament scale factors were 0.865, 1.000, 1.135, and 1.329 for the nominal D25, D50, D75, and D95 conditions. Only *E*_1_ (*ε*) through *E*_4_ (*ε*) were scaled:

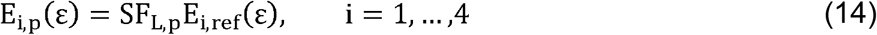

The equilibrium function *E*_∞_(ε) and all relaxation times remained unchanged.

**Table 2.** Ligament material-family assignments and reference strain-dependent constitutive functions.

| Material family | Ligament parts assigned to the family | Reference functions from Troyer and Puttlitz (2012) |
| --- | --- | --- |
| ALL-type | ALL; anterior atlanto-occipital ligament; apical ligament; posterior atlanto-occipital ligament; transverse atlantal ligament | $E_{\infty}(\varepsilon) = 18.38\varepsilon + 0.90$<br>$E_1(\varepsilon) = 14.70\varepsilon^2 + 0.24\varepsilon + 0.35$<br>$E_2(\varepsilon) = 3.28\varepsilon + 0.02$<br>$E_3(\varepsilon) = -2.16\varepsilon^2 + 3.51\varepsilon + 0.08$<br>$E_4(\varepsilon) = -7.95\varepsilon^2 + 5.02\varepsilon + 0.11$ |
| PLL-type | PLL; capsular ligament; tectorial membrane; alar ligament; interspinous ligament | $E_{\infty}(\varepsilon) = 9.79\varepsilon + 1.10$<br>$E_1(\varepsilon) = 1.13\varepsilon + 0.31$<br>$E_2(\varepsilon) = 1.09\varepsilon + 0.08$<br>$E_3(\varepsilon) = -0.30\varepsilon^2 + 1.23\varepsilon + 0.08$<br>$E_4(\varepsilon) = -5.89\varepsilon^2 + 2.51\varepsilon + 0.07$ |
| LF-type | Ligamentum flavum | $E_{\infty}(\varepsilon) = 1.20\varepsilon + 0.15$<br>$E_1(\varepsilon) = 2.03\varepsilon^2 - 0.17\varepsilon + 0.03$<br>$E_2(\varepsilon) = 0.53\varepsilon^2 + 0.07\varepsilon + 0.01$<br>$E_3(\varepsilon) = 1.17\varepsilon^2 - 0.17\varepsilon + 0.02$<br>$E_4(\varepsilon) = 0.68\varepsilon^2 + 0.04\varepsilon$ |

### 2.7 Intervertebral disc hyperviscoelasticity

The intervertebral discs were represented using the hyperviscoelastic formulation of Castro et al. (2014), as implemented in the previously developed head-neck model. The equilibrium hyperelastic response was represented with a Mooney-Rivlin strain-energy function:

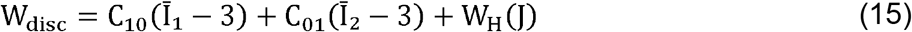

where *C*_10_ = 0.15*MPa, C*_01_ = 0.03 *MPa*, Ī_1_and Ī_2_are the first and second isochoric invariants, J is the volume ratio, and *W*_*H*_(*J*) represents the volumetric contribution. The transient response was represented using a three-term generalized Maxwell relaxation function:

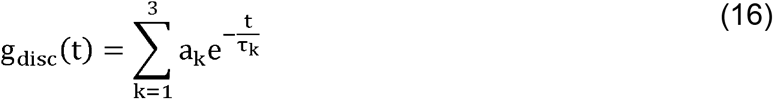

The reference coefficients were *a*_1_ = 1.7, *a*_2_ = 1.2, *and a*_3_ = 2.0, with relaxation times τ_1_ = 11.765 *s*, τ_2_ = 1.100*s*, and τ_3_ = 0.132*s* respectively (Castro et al., 2014).

Variability in disc viscoelasticity was estimated from the human intervertebral-disc damping measurements of Izambert et al. (2003). Coefficients of variation calculated from their reported mean and standard deviation values across loading frequencies had a median value of 0.33. The resulting disc scale factors were 0.777, 1.000, 1.223, and 1.543 for D25, D50, D75, and D95. Only the transient Maxwell coefficients were scaled 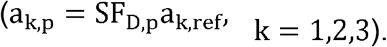.

The relaxation times, hyperelastic coefficients, and volumetric properties remained unchanged.

**Table 3.** Intervertebral-disc transient coefficients used in the passive-viscoelasticity conditions.

| Parameter | Nominal 25th | Reference | Nominal 75th | Nominal 95th |
| --- | --- | --- | --- | --- |
| $a_1$ | 1.321 | 1.700 | 2.079 | 2.623 |
| $a_2$ | 0.932 | 1.200 | 1.468 | 1.852 |
| $a_3$ | 1.554 | 2.000 | 2.446 | 3.086 |

### 2.8 Factorial simulation design and loading conditions

For the factorial analysis, S denotes muscle-strength level and D denotes passive cervical viscoelasticity level. The numbers 25, 50, 75, and 95 identify the four nominal levels used for each factor. Thus, for example, S25/D75 represents the S25 muscle-strength condition combined with the D75 passive-viscoelasticity condition. S50/D50 was used as the reference configuration. These labels identify literature-informed parameter levels and should not be interpreted as measured population percentiles for every individual muscle, ligament, or intervertebral-disc property. Muscle strength and passive cervical viscoelasticity were evaluated in a complete 4 × 4 factorial parameter grid. The strength levels S25, S50, S75, and S95 were combined with the passive-viscoelasticity levels D25, D50, D75, and D95, giving 16 cervical configurations per loading condition. For the strength conditions, the maximum isometric force of all cervical muscles was scaled by factors of 0.809, 1.000, 1.191, and 1.466 for S25, S50, S75, and S95, respectively. For the passive-viscoelasticity conditions, ligament and intervertebral-disc transient properties were varied together using their respective tissue-specific scaling factors. For example, D75 used a ligament scale factor of 1.135 and a disc scale factor of 1.223. The same 16 configurations were simulated under both loading conditions, resulting in 32 simulations overall.

Two mechanically distinct loading conditions were evaluated. The acceleration-driven condition was based on the frontal volunteer tests conducted at the Naval Biodynamics Laboratory (NBDL) by Ewing and Thomas (1972). A 15 g frontal acceleration condition was used, based on the corrected NBDL acceleration history reported by Thunnissen et al. (1995). The complete acceleration-time history was applied rather than a constant 15 g acceleration. This loading condition was implemented using the same approach previously used for validation of the baseline head-neck model (Bahreinizad et al., 2025).

The contact-driven condition represented a frontal American-football impact and was based on the loading approach of Mortensen et al. (2020). Mortensen et al. applied a 15-ms half-sine frontal force pulse with a peak force of 6800 N; their loading magnitude was based on reconstructed professional-football impacts reported by Viano et al. (2007), who reported impact forces of 7642 ± 2259 N. In the present study, the peak frontal contact force was set to 7000 N, providing a representative value within this reported range.

### 2.9 Numerical implementation

All simulations were solved using the LS-DYNA explicit solver. The brain ground-substance and axonal constitutive formulations were implemented as user-defined material models in Fortran. The embedded axonal network was coupled to the host brain mesh using the constrained beam-in-solid formulation.

Selective mass scaling was used to maintain a minimum stable time step of 1 × 10^−^□ s, with added mass limited to 5%. Model stability, total energy balance, hourglass energy, and added mass were checked for each simulation.

### 2.10 Atlas-based regional definition and outcome measures

Regional brain response was evaluated using the USCLobes atlas distributed through BrainSuite and described by Joshi et al. (2022). Fifteen regions were used: left and right frontal, parietal, temporal, occipital, insular, and cingulate regions, together with the brainstem, cerebellum, and corpus callosum. Atlas labels were registered to the subject-specific structural MRI and transferred to the FE brain coordinate system. Brain solid elements and axonal beam elements were assigned to the corresponding atlas region according to spatial location.

Global brain kinematics were evaluated at the brain center of mass (COM). Resultant linear acceleration and resultant angular acceleration were calculated from the three orthogonal components. Peak resultant brain COM linear and angular acceleration values were retained for comparison among parameter conditions.

Brain-skull relative motion was evaluated from the brain and skull centers of gravity. Three-dimensional brain displacement and skull displacement vectors were extracted separately. The skull displacement vector was subtracted from the brain displacement vector at each time point, and the resultant magnitude of this relative displacement vector was calculated. Peak resultant brain-skull relative displacement was reported for each simulation.

Regional axonal response was evaluated from the embedded tension-only beam elements. For each atlas region, axial strain values across all beam elements assigned to that region were ranked and the spatial 95th percentile was reported. This regional percentile was selected to reduce sensitivity to a small number of element-level outliers. Predicted axonal strain was interpreted as a mechanical measure of tract deformation and not as a direct clinical diagnosis of traumatic axonal injury.

Regional intracranial pressure was obtained from the brain solid elements. The absolute pressure values across all solid elements assigned to each atlas region were ranked, and the spatial 95th percentile was reported. This metric represents the short-duration dynamic pressure response during impact and should not be interpreted as sustained clinical intracranial hypertension.

## 3. Results

### 3.1 Effect of neck muscle strength

Greater neck muscle strength produced lower regional axonal strain throughout the brain under both loading conditions (Figs. 2 and 3). At the reference passive-viscoelasticity level (D50), the highest-strength condition (S95) produced lower 95th-percentile predicted axonal strain than the lowest-strength condition (S25) in all 15 brain regions. Under 15 g inertial loading, the S25-to-S95 reductions ranged from 10.30% in the brainstem to 58.13% in the left occipital region; the left and right frontal regions decreased by 22.07% and 23.34%, the left and right temporal regions by 24.36% and 25.57%, and the corpus callosum by 24.41%. Under 7000 N frontal loading, reductions ranged from 7.32% in the brainstem to 30.99% in the left occipital region, with decreases of 24.78% and 25.38% in the left and right temporal regions and 21.25% in the corpus callosum. Although S95 produced lower strain than S25 in every region, the response across the four strength levels was not monotonic in every region.

**Figure 2.**
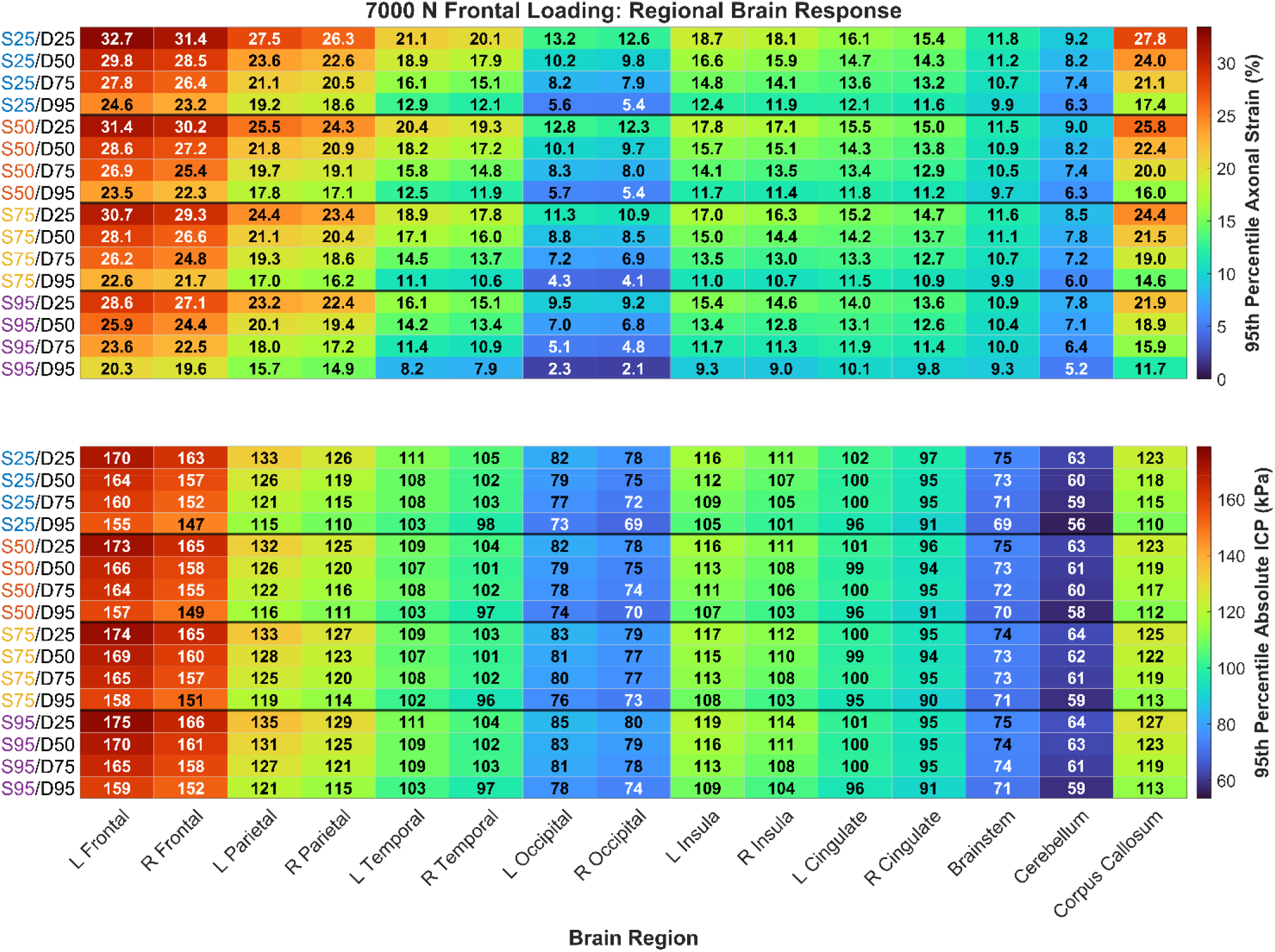
Regional 95th-percentile predicted axonal strain and absolute intracranial pressure across the 16 muscle-strength and passive-viscoelasticity combinations under 7000 N contact-driven frontal loading.

**Figure 3.**
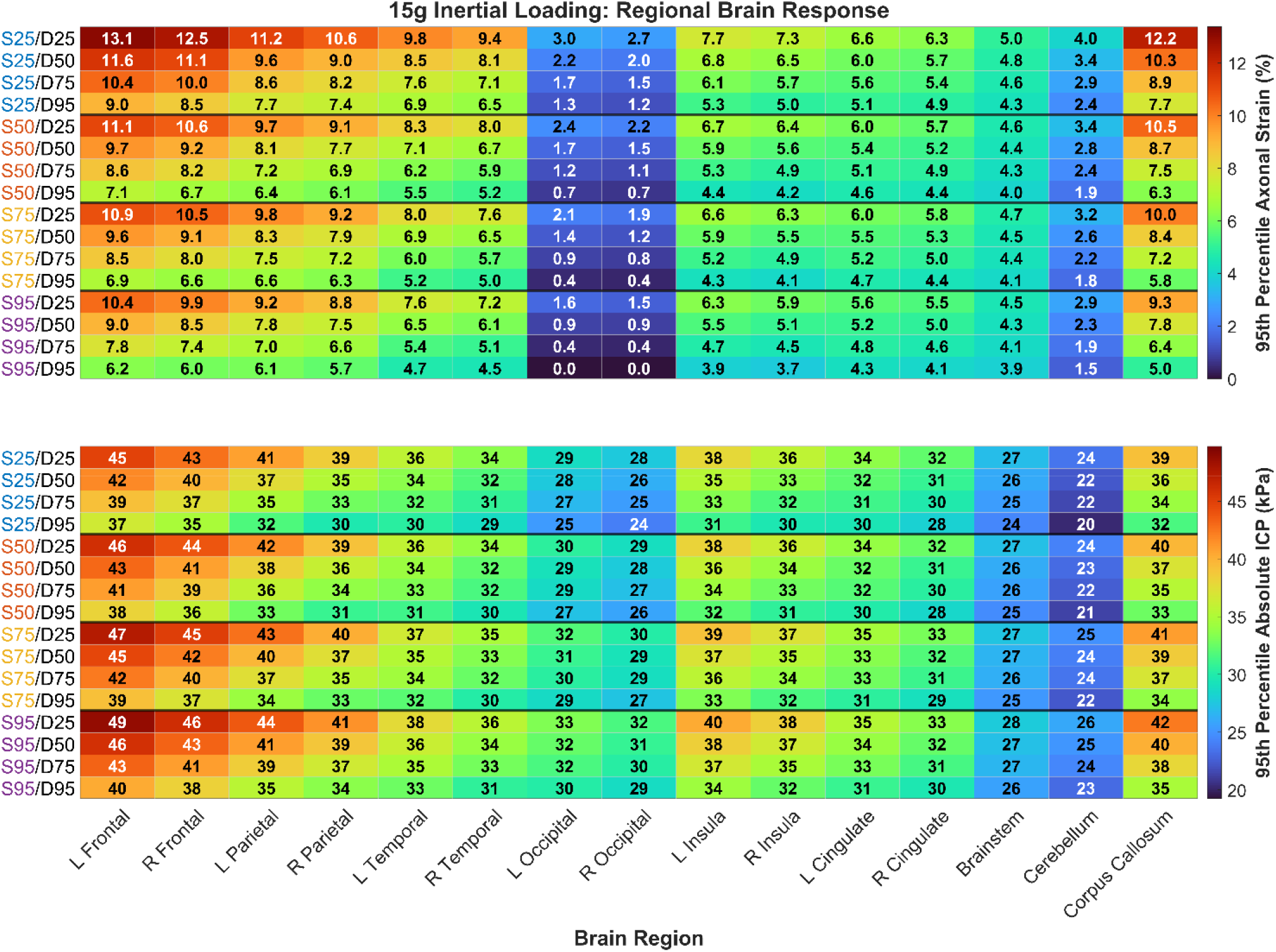
Regional 95th-percentile predicted axonal strain and absolute intracranial pressure across the 16 muscle-strength and passive-viscoelasticity combinations under 15 g acceleration-driven inertial loading.

The effect of muscle strength on global brain motion was small compared with its effect on regional axonal strain (Figs. 4 and 5). At D50, increasing strength from S25 to S95 changed brain-skull relative motion by −0.85% and −2.77%, brain COM linear acceleration by 0.00% and −0.67%, and brain COM angular acceleration by −1.18% and −2.17% under 15 g and 7000 N loading, respectively. None of the three global outcomes changed monotonically across all four strength levels. Muscle force-generating capacity therefore produced relatively small changes in peak global brain motion despite the larger changes in regional axonal strain.

**Figure 4.**
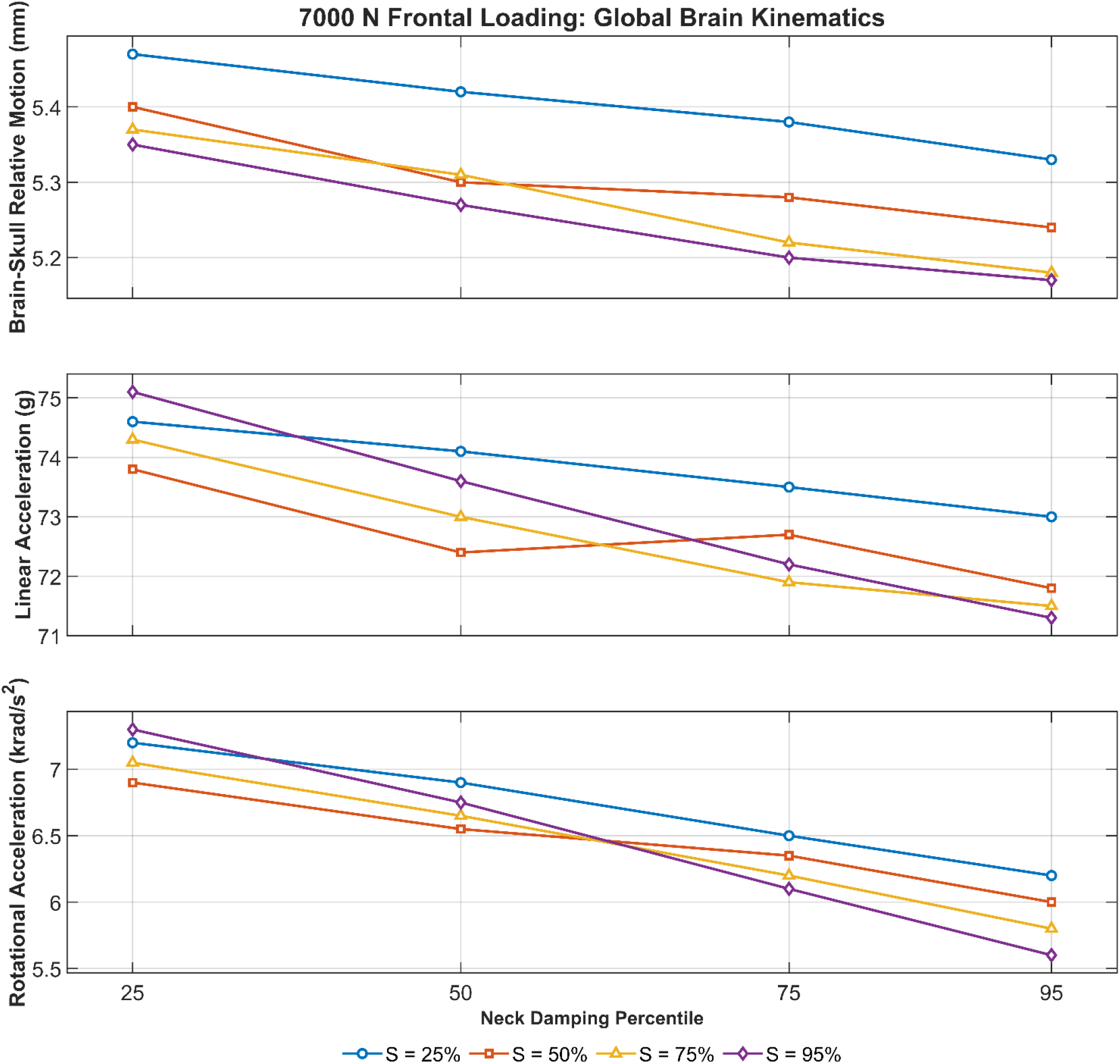
Brain-skull relative motion, brain COM linear acceleration, and brain COM angular acceleration across passive-viscoelasticity levels for the four muscle-strength levels under 7000 N frontal loading.

**Figure 5.**
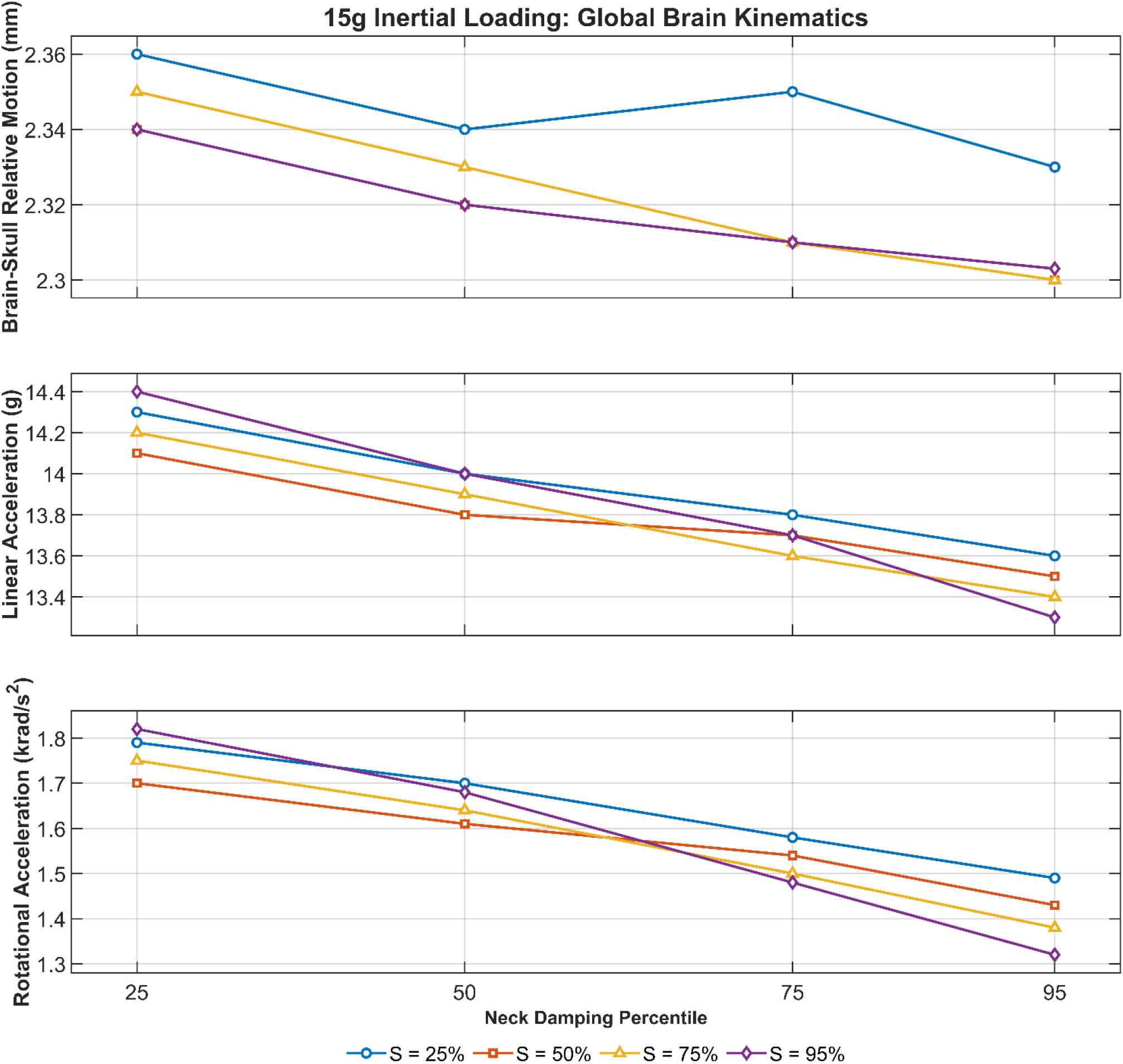
Brain-skull relative motion, brain COM linear acceleration, and brain COM angular acceleration across passive-viscoelasticity levels for the four muscle-strength levels under 15 g inertial loading.

Muscle strength affected regional 95th-percentile ICP magnitude differently from predicted axonal strain (Figs. 2 and 3). Under 15 g loading, increasing strength from S25 to S95 at D50 increased ICP magnitude in all 15 regions, with increases ranging from 4.06% in the brainstem to 17.29% in the left occipital region. Under 7000 N loading, the response was smaller and mixed, ranging from a 0.54% decrease in the right cingulate region to a 6.00% increase in the right occipital region. Thus, the lower predicted axonal strain at S95 was not accompanied by a corresponding reduction in ICP magnitude.

### 3.2 Effect of passive cervical viscoelasticity

Greater passive cervical viscoelasticity reduced regional predicted axonal strain under both loading conditions (Figs. 2 and 3). Under 15 g loading, the D25-to-D95 reductions ranged from 13.79% in the brainstem to 69.71% in the left occipital region; the left and right frontal regions decreased by 35.78% and 36.93%, the left and right temporal regions by 33.50% and 34.36%, the cerebellum by 43.63%, and the corpus callosum by 40.35%. Under 7000 N loading, reductions ranged from 15.75% in the brainstem to 56.06% in the right occipital region, with decreases of 38.58% and 38.61% in the left and right temporal regions and 37.93% in the corpus callosum. Unlike the muscle-strength response, the passive-viscoelasticity response was monotonic in all 15 regions under both loading conditions.

Passive cervical viscoelasticity also reduced global brain motion, with the largest relative change occurring in brain COM angular acceleration (Figs. 4 and 5). At S50, increasing passive cervical viscoelasticity from D25 to D95 reduced brain COM angular acceleration by 15.88% under 15 g loading and 13.04% under 7000 N loading. Brain COM linear acceleration decreased by 4.26% and 2.71%, while brain-skull relative motion decreased by 1.71% and 2.96%, respectively. The relative effect of passive cervical viscoelasticity was therefore substantially larger for angular acceleration than for linear acceleration or brain-skull relative motion.

Regional 95th-percentile ICP magnitude also decreased from D25 to D95 in every brain region under both loading conditions (Figs. 2 and 3). At S50, the reductions under 15 g loading ranged from 8.91% in the brainstem to 21.12% in the left parietal region. Under 7000 N loading, the reductions ranged from 5.07% in the left cingulate region to 11.89% in the left parietal region. In contrast to the muscle-strength response, greater passive cervical viscoelasticity reduced both predicted axonal strain and ICP magnitude.

### 3.3 Dependence between muscle strength and passive cervical viscoelasticity

The effect of muscle strength on regional predicted axonal strain increased as passive cervical viscoelasticity increased (Figs. 2 and 3). Using the arithmetic mean of the 15 regional 95th-percentile axonal-strain values as a descriptive summary, increasing strength from S25 to S95 reduced the response by 20.84%, 22.06%, 24.39%, and 28.33% at D25, D50, D75, and D95, respectively, under 15 g loading. Under 7000 N loading, the corresponding reductions were 17.38%, 17.55%, 19.25%, and 23.45%. The magnitude of the strength effect therefore increased across the passive-viscoelasticity levels under both loading conditions.

The effect of passive cervical viscoelasticity on regional predicted axonal strain also depended on muscle strength (Figs. 2 and 3). Increasing passive cervical viscoelasticity from D25 to D95 reduced the mean regional 95th-percentile axonal-strain response by 31.60%, 34.85%, 35.11%, and 38.07% at S25, S50, S75, and S95, respectively, under 15 g loading. Under 7000 N loading, the corresponding reductions were 32.79%, 32.52%, 33.60%, and 37.72%. The largest passive-viscoelasticity effect occurred at S95 under both loading conditions.

Brain COM angular acceleration showed the clearest dependence of the muscle-strength effect on passive cervical viscoelasticity (Figs. 4 and 5). Increasing strength from S25 to S95 changed angular acceleration by +1.68%, −1.18%, −6.33%, and −11.41% at D25, D50, D75, and D95, respectively, under 15 g loading and by +1.39%, −2.17%, −6.15%, and −9.68% under 7000 N loading. The direction of the strength effect therefore changed across passive-viscoelasticity levels: greater strength slightly increased angular acceleration at D25 but reduced it at D50, D75, and D95.

The effect of passive cervical viscoelasticity on brain COM angular acceleration also changed with muscle strength (Figs. 4 and 5). Increasing passive cervical viscoelasticity from D25 to D95 reduced angular acceleration by 16.76%, 15.88%, 21.14%, and 27.47% at S25, S50, S75, and S95, respectively, under 15 g loading. Under 7000 N loading, the corresponding reductions were 13.89%, 13.04%, 17.73%, and 23.29%. The largest reduction therefore occurred at S95 under both loading conditions.

Mean regional ICP changes varied less across levels of the other cervical factor than the corresponding axonal-strain and angular-acceleration changes (Figs. 2 and 3). Under 15 g loading, increasing strength from S25 to S95 increased the mean regional 95th-percentile ICP magnitude by 7.54%, 9.35%, 10.04%, and 10.22% at D25, D50, D75, and D95, respectively; under 7000 N loading, the corresponding changes were +1.63%, +2.91%, +3.18%, and +2.87%. Increasing passive cervical viscoelasticity from D25 to D95 reduced the mean regional ICP magnitude by 16.44%, 15.02%, 14.35%, and 14.36% at S25, S50, S75, and S95 under 15 g loading and by 9.36%, 8.30%, 7.97%, and 8.25% under 7000 N loading. The ICP response therefore showed less variation in factor-effect magnitude across the factorial grid than predicted axonal strain or angular acceleration.

Across the full S25/D25-to-S95/D95 range, regional predicted axonal strain and brain COM angular acceleration showed substantially larger relative changes than the other global motion outcomes (Figs. 2–5). The mean regional 95th-percentile axonal-strain response decreased by 50.98% under 15 g loading and 48.55% under 7000 N loading. Brain COM angular acceleration decreased by 26.26% and 22.22%, respectively, compared with reductions of 6.99% and 4.42% in brain COM linear acceleration and 2.42% and 5.48% in brain-skull relative motion.

Under 15 g loading, the occipital 95th-percentile predicted axonal-strain values approached zero at S95/D95 (Fig. 3). Left- and right-occipital strain decreased from 3.03% and 2.72% at S25/D25 to 0.03% and 0.04% at S95/D95, respectively. These absolute values are reported explicitly because the corresponding relative reductions were substantially larger than those observed in the other brain regions.

### 3.4 Brain-response validation

The updated axonal model produced higher correlations with the experimental brain-displacement histories than the baseline model in seven of the nine receiver-direction comparisons (Fig. 6). At Receiver 9, Pearson correlations for the axonal model were r = 0.68, 0.77, and 0.79 in the X, Y, and Z directions, respectively, compared with r = 0.70, 0.72, and 0.76 for the baseline model. At Receiver 16, the corresponding axonal-model correlations were r = 0.40, 0.94, and 0.71, compared with r = 0.41, 0.91, and 0.57 for the baseline model. At Receiver 31, correlations were r = 0.46, 0.82, and 0.92 for the axonal model and r = 0.41, 0.80, and 0.90 for the baseline model. Overall, correlations ranged from 0.40 to 0.94 for the updated axonal model and from 0.41 to 0.91 for the baseline model. The largest improvement occurred in the Z direction at Receiver 16, while small decreases were observed in the X direction at Receivers 9 and 16.

**Figure 6.**
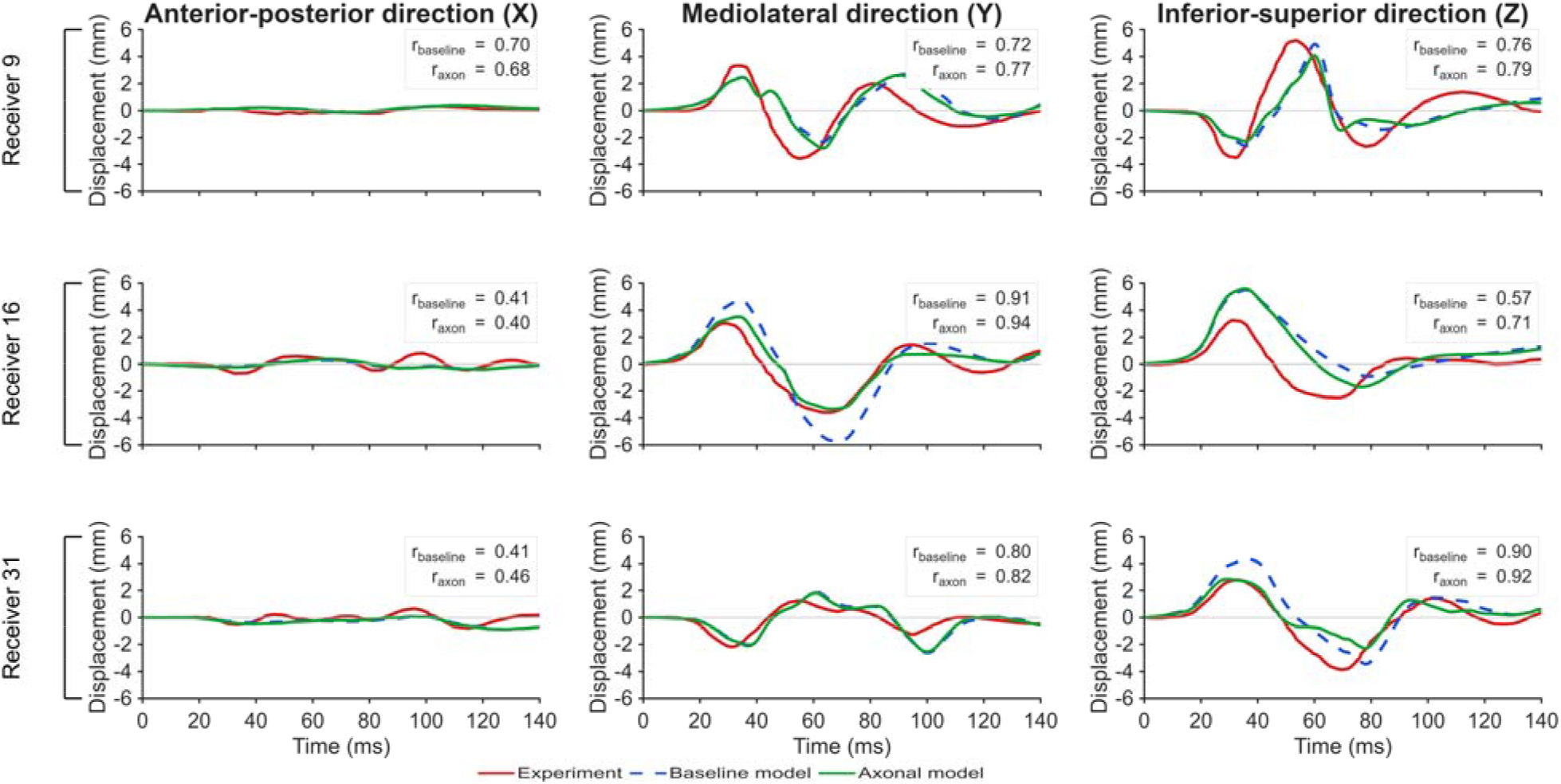
Brain-displacement validation against Alshareef et al. (2018), comparing the experimental response with the baseline and updated axonal models at Receivers 9, 16, and 31. Pearson correlation coefficients are reported for each receiver and displacement direction.

## 4. Discussion

### 4.1 Main findings

The main finding of this study is that passive cervical viscoelasticity produced a larger change in regional brain response than muscle force-generating capacity across the tested parameter ranges. At the reference level of the other factor, increasing passive viscoelasticity reduced the mean regional axonal-strain metric by 34.9% under inertial loading and 32.5% under frontal loading, compared with 22.1% and 17.5% reductions from increasing muscle strength. Passive viscoelasticity also reduced brain COM angular acceleration by 15.9% and 13.0%, whereas the corresponding strength effects were 1.2% and 2.2%. The two factors therefore influenced the same head-brain system through different mechanical pathways and with different magnitudes.

The second major finding is that global kinematic changes were much smaller than many regional axonal-strain changes. Strength variation changed relative motion, linear acceleration, and angular acceleration by only a few percent at D50, while regional axonal-strain reductions reached 58.1% under inertial loading and 31.0% under frontal loading. Passive-viscoelasticity variation showed the same separation, with small relative-motion changes and regional strain reductions as large as 69.7% and 56.1%. The local brain response therefore cannot be inferred from a single global peak metric.

The third major finding is that pressure and axonal strain did not respond in parallel. Greater passive viscoelasticity reduced both outcomes, while greater muscle strength reduced axonal strain and increased dynamic ICP under inertial loading. This divergence shows that a cervical change can redistribute the mechanical response rather than uniformly lowering every intracranial metric.

### 4.2 Why passive cervical viscoelasticity produced the larger effect

Passive cervical tissues had a larger effect because ligaments and discs resist rapid motion in a time-dependent manner. Troyer and Puttlitz (2011, 2012) showed that human cervical ligaments require a nonlinear viscoelastic description, and Lucas et al. (2008) measured substantial fast-rate time dependence in major cervical ligaments. In the present ligament formulation, increasing the transient terms raises the short-time resistance generated before relaxation develops, while the equilibrium term and relaxation times remain unchanged. The disc formulation changes the same part of the response by scaling the transient Maxwell coefficients while preserving the equilibrium Mooney-Rivlin parameters. Izambert et al. (2003) independently demonstrated frequency-dependent dynamic stiffness and damping in human intervertebral discs. These constitutive features make passive cervical resistance especially relevant during rapid head loading.

The current results also agree with the head-impact mechanism reported by Kuo et al. (2019). Their OpenSim analysis showed that active musculature and passive ligaments contribute differently as impact severity and loading rate change, with rate-dependent passive ligaments becoming relatively more important during high-rate head motion. In the present FE model, passive-viscoelasticity variation changed brain COM angular acceleration far more than strength variation under both loading conditions. The agreement is mechanistically important: the current study extends the role of passive cervical tissues from head stabilization to regional brain and axonal response.

### 4.3 Muscle strength changed local deformation with modest global kinematic change

The direction of the muscle-strength effect is consistent with controlled experimental and computational studies, while its magnitude reinforces the limits of a simple stronger-neck interpretation. Eckner et al. (2014) found inverse relationships between isometric neck strength and impulsive head linear and angular velocity, with correlation coefficients of 0.42–0.66. Mortensen et al. (2020) similarly found that increasing cervical muscle strength reduced modeled head-injury metrics, although posture and impact characteristics had larger effects. Jin et al. (2017) also found that active cervical muscle responses could reduce several predicted brain-injury measures and that anticipatory activation played a dominant role. In contrast, Schmidt et al. (2014) found no reduction in football head-impact severity from greater muscle strength or muscle size, and Garrett et al. (2023) reported only small nonsignificant pooled relationships between greater neck strength and concussion risk. The present results fit this mixed literature: strength modified the mechanical response, although it did not strongly reduce global acceleration.

Holding activation fixed is important for interpreting the present strength effect. The model increased Fmax while preserving the same OpenSim-derived activation pattern across all cases. The simulations therefore isolate available force capacity rather than anticipation, reflex timing, or voluntary cocontraction. Jin et al. (2017) showed that anticipatory activation can dominate the response in an impact simulation, so the current results should not be interpreted as the total effect of the neuromuscular system. They specifically represent the effect of changing force-generating capacity within the same activation strategy.

### 4.4 Dependence Between Muscle Strength and Passive Cervical Viscoelasticity

The effect of muscle strength depended on the passive mechanical state of the neck, and the effect of passive viscoelasticity depended on muscle strength. The strength-associated reduction in the mean regional axonal-strain metric increased from 20.8% at D25 to 28.3% at D95 under inertial loading and from 17.4% to 23.4% under frontal loading. The passive-viscoelasticity effect also increased at the highest strength level. This pattern was strongest in angular acceleration, where increasing strength slightly increased the peak at D25 and reduced it substantially at D95. The direction and magnitude of the muscle-strength effect therefore depended on the passive mechanical state of the neck.

This conditional behavior is consistent with a coupled dynamic system rather than a set of independent linear modifiers. Muscle force, ligament resistance, disc response, head inertia, and external loading act simultaneously through the cervical degrees of freedom. A change in one component alters the motion state seen by the others, so the final response can differ from the sum of isolated one-factor effects. The deterministic design does not provide a population-level interaction test, although the full grid clearly shows that effect magnitude changes across factor combinations.

### 4.5 Regional strain, brain-skull motion, and pressure represent different mechanics

The large regional strain changes relative to the small brain-skull relative-motion changes are consistent with previous validation research. Zhou et al. (2018) reported no significant correlation between peak brain strain and peak brain-skull relative motion and also found that validation scores for the two response types were not significantly correlated. The present results show the same separation within one model: passive-viscoelasticity variation reduced relative motion by only 1.7-3.0% at S50 while reducing the mean regional axonal-strain metric by 32.5-34.9%. Brain-skull displacement is therefore a useful global motion measure, although it cannot serve as a direct surrogate for local tract deformation.

The different pressure and strain trends are also consistent with established head-impact mechanics. In accident reconstructions, Kleiven (2007) found that translational kinematics contributed little to distortional brain strain, while rotational kinematics contributed little to the pressure response. The present muscle-strength cases showed the same type of decoupling: axonal strain decreased throughout the brain while dynamic absolute ICP increased under inertial loading. Pressure and axonal strain should therefore be interpreted as complementary rather than interchangeable outcomes.

Regional tract orientation and anatomy provide a second reason for the nonuniform strain response. Giordano and Kleiven (2014), Ji et al. (2015), and Zhao et al. (2016) showed that strain resolved along white-matter pathways differs from conventional tissue-level strain in both magnitude and spatial distribution. Wu et al. (2019) and Hajiaghamemar et al. (2020) further showed the value of explicit tract representations for evaluating axonal mechanics. In the present model, the same change in head-neck dynamics produced different strain changes across frontal, temporal, occipital, callosal, cerebellar, and brainstem regions. The neck therefore changed the spatial distribution of tract deformation rather than uniformly scaling the whole brain.

### 4.6 Clinical relevance of cervical mechanical variability

Among the cervical factors evaluated in this study, muscle strength is the one that can be modified most directly through training. Catenaccio et al. (2017) reported cervical strength values ranging from 38 to 383 N in men and from 15 to 223 N in women, demonstrating large differences in force-generating capacity among healthy individuals. Neck strength can also be increased through resistance training (Geary et al., 2014). However, greater strength has not consistently translated to lower head-impact severity or concussion risk. Eckner et al. (2014) reported smaller changes in head linear and angular velocity in participants with greater isometric neck strength during controlled impulsive loading, whereas Mihalik et al. (2011) and Schmidt et al. (2014) did not find a consistent reduction in head-impact severity with greater cervical strength. Garrett et al. (2023) similarly found no significant pooled association between greater neck strength and lower concussion risk. Evidence is even more limited for post-concussion recovery, and it remains unclear whether greater preinjury neck strength reduces symptom severity or shortens recovery time. The present simulations therefore show that muscle force-generating capacity can alter brain mechanics, but they do not establish that neck strengthening prevents concussion or improves clinical recovery.

The cervical spine may nevertheless be important after concussion. Neck pain, stiffness, and restricted cervical motion are common in patients with concussion and can contribute to headache, dizziness, and other persistent symptoms. Oyekan et al. (2023) reported that patients with more severe neck symptoms had greater overall symptom burden and required approximately 40 days to recover, compared with approximately 30 days in patients without severe neck symptoms. Cervical rehabilitation has also shown benefit in selected patients with persistent symptoms. Schneider et al. (2014), for example, found that patients receiving combined cervical and vestibular rehabilitation were more likely to be medically cleared within eight weeks than those receiving standard care alone. These clinical findings do not show that the mechanical changes predicted in the present model directly determine recovery, but they demonstrate that cervical function can remain clinically relevant after the initial head impact.

Passive cervical mechanics may provide another source of individual variability. Reduced cervical mobility can occur after cervical fusion because one or more motion segments are mechanically constrained. This does not mean that cervical fusion is equivalent to the lower passive-viscoelasticity conditions used in the present model, but it illustrates that the mechanical boundary condition between the head and torso can differ substantially among individuals. At the opposite end of the mobility spectrum, Ehlers-Danlos syndrome can involve ligamentous laxity, cervical hypermobility, and craniocervical instability (Mao et al., 2022). Limited clinical evidence also suggests that some patients with Ehlers-Danlos syndrome may experience slower or less complete recovery following mild traumatic brain injury (Gami & Singman, 2019). These observations do not establish a direct relationship between cervical viscoelasticity and concussion outcome, but they provide clinically relevant examples of conditions in which cervical mechanical behavior differs from that of a typical healthy neck.

The present results should also not be interpreted as evidence that flexibility exercises can directly modify the passive-viscoelasticity parameter evaluated here. Cervical rehabilitation can improve range of motion, pain, neuromuscular control, and functional mobility, but these outcomes are different from changing the intrinsic time-dependent material properties of cervical ligaments and intervertebral discs. Whether those passive tissue properties can be meaningfully modified through rehabilitation, and whether such changes would alter brain mechanics during head impact, remains unknown. The current findings therefore support further investigation of cervical mechanical variability as a possible contributor to differences in brain response, rather than a specific recommendation for neck strengthening or flexibility training.

### 4.7 Interpretation of predicted axonal strain

Predicted axonal strain is a mechanical surrogate and not a direct diagnosis of traumatic axonal injury. Bain and Meaney (2000) demonstrated strain-dependent morphological and electrophysiological injury in dynamically stretched white matter, and Hajiaghamemar and Margulies (2021) showed that an embedded white-matter tract model could predict the location of traumatic diffuse axonal injury in an animal model. These studies support the biological relevance of tract-oriented deformation, although their injury thresholds cannot be transferred directly to the current human FE model.

The embedded-tract approach nevertheless provides an important modeling advantage for this study. Wu et al. (2019) represented white matter as an isotropic ground substance with explicitly embedded axonal fiber tracts, allowing deformation to be calculated directly along the tract structures. The current model applies that framework to subject-specific DTI-derived pathways within a full head-neck model. The resulting axonal strain values are best used for controlled comparisons among cervical conditions rather than as absolute clinical injury thresholds.

### 4.8 Limitations

This study has several limitations that define the scope of the conclusions. First, the model represents one male anatomy, so geometry remained fixed while selected muscle, ligament, and disc properties were varied. Second, muscle strength was scaled uniformly across all 42 muscles. Real individuals do not change every muscle by the same factor, and muscle-specific architecture, moment arms, hypertrophy, and neural recruitment vary across people. Uniform scaling was used intentionally to isolate whole-neck force capacity. Muscle activation was also held constant, so the study does not represent differences in anticipation or reflex response. Third, the passive-viscoelasticity levels are literature-informed sensitivity conditions rather than measured cervical population percentiles. Ligament variability was derived from the fast-rate experimental data of Lucas et al. (2008), while disc variability was estimated from lumbar-disc damping measurements reported by Izambert et al. (2003). Fourth, ligament and disc transient properties were varied together as one passive-cervical factor. The current design therefore quantifies their combined contribution and does not separate the independent effects of discs, lower-cervical ligaments, and upper-cervical ligament structures. Tissue-specific studies would be needed to resolve those contributions. Fifth, only two loading conditions were evaluated. Kuo et al. (2019) showed that cervical muscle and ligament contributions depend on loading severity and direction, so the present trends cannot be assumed to apply unchanged to lateral, oblique, extension, axial-rotation, or longer-duration impacts.

## 5. Conclusion

Neck muscle strength and passive cervical viscoelasticity both changed predicted brain response, although passive cervical viscoelasticity produced the larger effect across the tested ranges. At the reference level of the other factor, increasing strength reduced the mean regional axonal-strain metric by 22.1% under 15 g inertial loading and 17.5% under 7000 N frontal loading, while increasing passive viscoelasticity reduced the same metric by 34.9% and 32.5%. Passive viscoelasticity also reduced brain COM angular acceleration by 15.9% and 13.0%, compared with 1.2% and 2.2% reductions from strength.

The two cervical factors also changed intracranial outcomes differently. Greater passive viscoelasticity reduced regional dynamic ICP and predicted axonal strain, while greater muscle strength reduced axonal strain and increased ICP under inertial loading. The full 4 × 4 design further showed that the magnitude of the strength effect increased as passive viscoelasticity increased, with the clearest conditional behavior in angular acceleration. The neck therefore acts as a coupled mechanical boundary condition rather than as a single strength parameter.

These findings support including cervical mechanical variability in future subject-specific TBI models. Muscle force capacity is one modifiable component, while passive ligament and disc mechanics represent a separate source of biological and clinical variability. The external head load alone does not determine regional brain deformation; the mechanical state of the neck changes how that load reaches the brain.

## References

Alshareef, A., Giudice, J. S., Forman, J., Salzar, R. S., & Panzer, M. B. (2018). A novel method for quantifying human in situ whole brain deformation under rotational loading using sonomicrometry. Journal of Neurotrauma, 35, 780–789.

Alvarez, V. S., Halldin, P., & Kleiven, S. (2014). The influence of neck muscle tonus and posture on brain tissue strain in pedestrian head impacts. Stapp Car Crash Journal, 58, 63–101.

Bahreinizad, H., Paulon, G. M., Wei, L., & Chowdhury, S. K. (2025). An MRI-derived head-neck finite element model. Biomechanics and Modeling in Mechanobiology, 24, 2195– 2215.

Bain, A. C., & Meaney, D. F. (2000). Tissue-level thresholds for axonal damage in an experimental model of central nervous system white matter injury. Journal of Biomechanical Engineering, 122, 615–622.

Castro, A. P. G., Paul, C. P. L., Detiger, S. E. L., Smit, T. H., van Royen, B. J., Pimenta Claro, J. C., Mullender, M. G., & Alves, J. L. (2014). Long-term creep behavior of the intervertebral disk: Comparison between bioreactor data and numerical results. Frontiers in Bioengineering and Biotechnology, 2, 56.

Catenaccio, E., Mu, W., Kaplan, A., Fleysher, R., Kim, N., Bachrach, T., Zughaft Sears, M., Jaspan, O., Caccese, J., Kim, M., Wagshul, M., Stewart, W. F., Lipton, R. B., & Lipton, M. L. (2017). Characterization of neck strength in healthy young adults. PM&R, 9, 884–891.

Cournoyer, J., Koncan, D., Gilchrist, M. D., & Hoshizaki, T. B. (2021). The influence of neck stiffness on head kinematics and maximum principal strain associated with youth American football collisions. Journal of Applied Biomechanics, 37, 288–295.

Dewan, M. C., Rattani, A., Gupta, S., Baticulon, R. E., Hung, Y. C., Punchak, M., Agrawal, A., Adeleye, A. O., Shrime, M. G., Rubiano, A. M., Rosenfeld, J. V., & Park, K. B. (2019). Estimating the global incidence of traumatic brain injury. Journal of Neurosurgery, 130, 1080–1097.

Dirisala, V., Karami, G., & Ziejewski, M. (2012). Effects of neck damping properties on brain response underimpact loading. International Journal for Numerical Methods in Biomedical Engineering, 28, 472–494.

Eckner, J. T., Oh, Y. K., Joshi, M. S., Richardson, J. K., & Ashton-Miller, J. A. (2014). Effect of neck muscle strength and anticipatory cervical muscle activation on the kinematic response of the head to impulsive loads. American Journal of Sports Medicine, 42, 566– 576.

Ewing, C. L., & Thomas, D. J. (1972). Human head and neck response to impact acceleration. NAMRL Monograph 21; USAARL 73-1. Naval Aerospace Medical Research Laboratory.

Farber, S. H., Mugge, L. A., Sawa, A. G. U., DiDomenico, J. D., Assefa, T. G., Ratliff, A. C., Crawford, N. R., Uribe, J. S., Kelly, B. P., & Turner, J. D. (2025). Clinical effect of occipitocervical and subaxial cervical fusion constructs on range of motion: Comprehensive guide based on biomechanical cadaveric testing on 1009 motion segments. Journal of Neurosurgery: Spine, 43, 352–360.

Gami, A., & Singman, E. L. (2019). Underlying Ehlers-Danlos syndrome discovered during neuro-ophthalmic evaluation of concussion patients: A case series. BMC Ophthalmology, 19, 159.

Garrett, J. M., Mastrorocco, M., Peek, K., van den Hoek, D. J., & McGuckian, T. B. (2023). The relationship between neck strength and sports-related concussion in team sports: A systematic review with meta-analysis. Journal of Orthopaedic & Sports Physical Therapy, 53, 585–593.

Geary, K., Green, B. S., & Delahunt, E. (2014). Effects of neck strength training on isometric neck strength in rugby union players. Clinical Journal of Sport Medicine, 24, 502– 508.

Giordano, C., & Kleiven, S. (2014). Evaluation of axonal strain as a predictor for mild traumatic brain injuries using finite element modeling. Stapp Car Crash Journal, 58, 29–61.

Hajiaghamemar, M., Wu, T., Panzer, M. B., & Margulies, S. S. (2020). Embedded axonal fiber tracts improve finite element model predictions of traumatic brain injury. Biomechanics and Modeling in Mechanobiology, 19, 1109–1130.

Hajiaghamemar, M., & Margulies, S. S. (2021). Multi-scale white matter tract embedded brain finite element model predicts the location of traumatic diffuse axonal injury. Journal of Neurotrauma, 38, 144–157.

Ito, S., Ivancic, P. C., Pearson, A., Tominaga, Y., Gimenez, S., Rubin, W., & Panjabi, M. M. (2005). Cervical intervertebral disc injury during simulated frontal impact. European Spine Journal, 14, 356–365.

Izambert, O., Mitton, D., Thourot, M., & Lavaste, F. (2003). Dynamic stiffness and damping of human intervertebral disc using axial oscillatory displacement under a free mass system. European Spine Journal, 12, 562–566.

Ji, S., Zhao, W., Ford, J. C., Beckwith, J. G., Bolander, R. P., Greenwald, R. M., Flashman, L. A., Paulsen, K. D., & McAllister, T. W. (2015). Group-wise evaluation and comparison of white matter fiber strain and maximum principal strain in sports-related concussion. Journal of Neurotrauma, 32, 441–454.

Jin, X., Feng, Z., Mika, V., Li, H., Viano, D. C., & Yang, K. H. (2017). The role of neck muscle activities on the risk of mild traumatic brain injury in American football. Journal of Biomechanical Engineering, 139, 101002.

Joshi, A. A., Choi, S., Liu, Y., Chong, M., Sonkar, G., Gonzalez-Martinez, J., Nair, D., Wisnowski, J. L., Haldar, J. P., Shattuck, D. W., Damasio, H., & Leahy, R. M. (2022). A hybrid high-resolution anatomical MRI atlas with sub-parcellation of cortical gyri using resting fMRI. Journal of Neuroscience Methods, 374, 109566.

Kleiven, S. (2007). Predictors for traumatic brain injuries evaluated through accident reconstructions. Stapp Car Crash Journal, 51, 81–114.

Kuo, C., Sheffels, J., Fanton, M., Yu, I. B., Hamalainen, R., & Camarillo, D. (2019). Passive cervical spine ligaments provide stability during head impacts. Journal of the Royal Society Interface, 16, 20190086.

Lucas, S. R., Bass, C. R., Salzar, R. S., Oyen, M. L., Planchak, C., Ziemba, A., Shender, B. S., & Paskoff, G. (2008). Viscoelastic properties of the cervical spinal ligaments under fast strain-rate deformations. Acta Biomaterialia, 4, 117–125.

Mao, G., Kopparapu, S., Jin, Y., Davidar, A. D., Hersh, A. M., Weber-Levine, C., & Theodore, N. (2022). Craniocervical instability in patients with Ehlers-Danlos syndrome: Controversies in diagnosis and management. The Spine Journal, 22, 1944–1952.

Mihalik, J. P., Guskiewicz, K. M., Marshall, S. W., Greenwald, R. M., Blackburn, J. T., & Cantu, R. C. (2011). Does cervical muscle strength in youth ice hockey players affect head impact biomechanics? Clinical Journal of Sport Medicine, 21, 416–421.

Mortensen, J. D., Vasavada, A. N., & Merryweather, A. S. (2018). The inclusion of hyoid muscles improve moment generating capacity and dynamic simulations in musculoskeletal models of the head and neck. PLOS ONE, 13, e0199912.

Mortensen, J. D., Vasavada, A. N., & Merryweather, A. S. (2020). Sensitivity analysis of muscle properties and impact parameters on head injury risk in American football. Journal of Biomechanics, 100, 109411.

Oyekan, A. A., Eagle, S., Trbovich, A. M., Shaw, J. D., Schneider, M., Collins, M., Lee, J. Y., & Kontos, A. P. (2023). Neck symptoms and associated clinical outcomes in patients following concussion. Journal of Head Trauma Rehabilitation, 38, 417–424.

Panzer, M. B., Fice, J. B., & Cronin, D. S. (2011). Cervical spine response in frontal crash. Medical Engineering & Physics, 33, 1147–1159.

Schmidt, J. D., Guskiewicz, K. M., Blackburn, J. T., Mihalik, J. P., Siegmund, G. P., & Marshall, S. W. (2014). The influence of cervical muscle characteristics on head impact biomechanics in football. American Journal of Sports Medicine, 42, 2056–2066.

Schneider, K. J., Meeuwisse, W. H., Nettel-Aguirre, A., Barlow, K., Boyd, L., Kang, J., & Emery, C. A. (2014). Cervicovestibular rehabilitation in sport-related concussion: A randomised controlled trial. British Journal of Sports Medicine, 48, 1294–1298.

Thunnissen, J., Wismans, J., Ewing, C., & Thomas, D. (1995). Human volunteer head-neck response in frontal flexion: A new analysis. SAE Transactions, 104, 3065–3086.

Troyer, K. L., & Puttlitz, C. M. (2011). Human cervical spine ligaments exhibit fully nonlinear viscoelastic behavior. Acta Biomaterialia, 7, 700–709.

Troyer, K. L., & Puttlitz, C. M. (2012). Nonlinear viscoelasticity plays an essential role in the functional behavior of spinal ligaments. Journal of Biomechanics, 45, 684–691.

Vasavada, A. N., Danaraj, J., & Siegmund, G. P. (2008). Head and neck anthropometry, vertebral geometry and neck strength in height-matched men and women. Journal of Biomechanics, 41, 114–121.

Viano, D. C., Casson, I. R., & Pellman, E. J. (2007). Concussion in professional football: Biomechanics of the struck player—Part 14. Neurosurgery, 61(2), 313–327.

Wu, T., Alshareef, A., Giudice, J. S., & Panzer, M. B. (2019). Explicit modeling of white matter axonal fiber tracts in a finite element brain model. Annals of Biomedical Engineering, 47, 1908–1922.

Zhao, W., Ford, J. C., Flashman, L. A., McAllister, T. W., & Ji, S. (2016). White matter injury susceptibility via fiber strain evaluation using whole-brain tractography. Journal of Neurotrauma, 33, 1834–1847.

Zhou, Z., Li, X., Kleiven, S., Shah, C. S., & Hardy, W. N. (2018). A reanalysis of experimental brain strain data: Implication for finite element head model validation. Stapp Car Crash Journal, 62, 293–318.

